# Evaluating clinical stop-smoking services globally: Proposal for a minimum data set

**DOI:** 10.1101/126078

**Authors:** Andrew L. Skinner, Robert West, Martin Raw, Emma Anderson, Marcus R. Munafò

## Abstract

**Background and aims:** Behavioural and pharmacological support for smoking cessation improves the chances of success and represents a highly cost-effective way of preventing chronic disease and premature death. There are a large number of clinical stop-smoking services around the world. These could be connected into a global network to provide data to assess what treatment components are most effective, for what populations, in what settings. This requires data to be collected according to a minimum standard set of data items. This paper sets out a proposal for this global minimum data set.

**Methods:** We reviewed sets of data items used in clinical services that have already benefited from standardised approaches to using data. We identified client and treatment data items that may directly or indirectly influence outcome, and outcome variables that were practicable to obtain in clinical practice. We then consulted service providers in countries that may have an interest in taking part in a global network of smoking cessation services, and revised the sets of data items according to their feedback.

**Results:** Three sets of data items are proposed. The first is a set of features characterising treatments offered by a service. The second is a core set of data items describing clients’ characteristics, engagement with the service, and outcomes. The third is an extended set of client data items to be captured in addition to the core data items wherever resources permit.

**Conclusions:** We propose minimum standards for capturing data from clinical smoking cessation services globally. This could provide a basis for meaningful evaluations of different smoking cessation treatments in different populations in a variety of settings across many countries.

## Introduction

There are currently over 1 billion tobacco users worldwide [1]. Despite the prevalence of tobacco use decreasing in many countries, this number is not falling, partly because of increasing prevalence in other countries and partly because of population growth. Some of these users may be able to stop when they choose, but the success rates of unaided quit attempts [2] and findings from clinical trials and population studies [3] indicate that most would benefit from support in their attempt to stop smoking. Article 14 of the World Health Organization Framework Convention on Tobacco Control (FCTC) requires Parties to take effective measures to promote tobacco use cessation and adequate treatment for tobacco dependence. Guidelines for the implementation of Article 14 were adopted in November 2010 and outline what is needed to enable Parties to meet their obligations under that Article [4]. Current evidence, however, indicates that only a small minority of countries, and very few low- and middle-income countries, have the infrastructure and systems elements in places to be able to meet these obligations [5, 6].

Studies in some high-income countries suggest specific medicines and types of behavioural support can improve smokers’ chances of long-term success at stopping [7]. However, in order to apply this work globally we ideally need to know more about real world effectiveness throughout the world. More broadly, there is considerable room for improvement in stop-smoking support in all settings. We need to build incrementally on our understanding of how combinations of behaviour change techniques, delivered in what way, to what kinds of smoker in what settings provide the optimal outcomes [7].

In the future, the challenge for tobacco use cessation will be to improve on the strategies currently available and more importantly, to identify support options that are clearly effective, cost effective, and affordable in different populations and different regions. Randomised controlled trials (RCTs) can only take us so far in this endeavour, and suffer from the well-established limitations of generalizability, practicability, cost and timescale [8].

A programme of research is required, complementing the existing evidence base from RCTs, which provides vital information about what approaches work best across a broad range of settings. This programme should consist of quasi-experimental and epidemiological studies that examine variations in smoking cessation outcomes among different cohorts of smokers using different methods of quitting, adjusting for as many possible confounding factors as possible [9]. This kind of approach has already paid dividends in the United Kingdom, where the combination of a common national standard for outcome assessment [10], and widespread use of a single database for recording crucial information, has enabled confirmation of findings from RCTs concerning the relative effectiveness of different forms of medication, and identification of specific treatments with improved success rates [11].

Our vision is to extend the success of this approach globally. By connecting clinical smoking cessation services from many countries to form a global network of service providers, it will be possible to share information about the performance of different treatments in different scenarios within a common frame of reference, and to assess and identify optimal treatment strategies for different populations in different settings on a global scale. A key enabler in connecting providers to form a global network will be establishing a standardised, minimum set of data that should be captured by all clinical smoking cessation services in the network. Here, we take the first step towards defining this minimum data set.

We propose data sets that describe the treatments offered by services, and client’s engagement with these services. The data required to describe client’s engagement with smoking cessation services can be extensive. Regional differences in resources (e.g., computer equipment, network infrastructure, measurement devices) means the extent of data that can be captured by services will vary. We therefore suggest that, rather than a single client data set, we define a core set of that should be captured in every smoking cessation service setting, and an extended, richer set of data items that should be captured wherever resources make this possible. We propose candidate sets of core and extended data items.

Not all countries have sufficient infrastructure to support dedicated clinical smoking cessation services. In those that do not, there is often more of a focus on delivery of brief advice supporting smoking cessation through alternative, existing mechanisms, such as tuberculosis clinics [12, 13]. These brief advice approaches vary considerably from the services offered by smoking cessation clinics, largely because they are not designed to follow-up and track client quit attempts. Consequently, the data describing brief advice approaches will vary significantly from data describing clinical smoking cessation services. We therefore propose that defining minimum data sets for brief advice approaches should be a separate (but linked) activity, and do not address this here.

## Methods

For the treatment data, we took as our starting point a list of behaviour change techniques found in UK Stop Smoking Services (SSS) [14]. From this we selected only those treatment features found to be associated with improved quit outcomes (either self-reported or CO verified).

For client data, we began with the data items listed in the latest version (v9) of the United Kingdom’s National Centre for Smoking Cessation and Training’s Stop Smoking Service Client Record Form [15]. Through an iterative process of modification and review, the authors, who have extensive experience of international smoking cessation services and epidemiological studies of tobacco use, arrived at initial sets of core and extended client data items. As this process of selection was more organic, justification for the inclusion of each client data item is included in Tables 2 and 3.

**Table 1 –.**
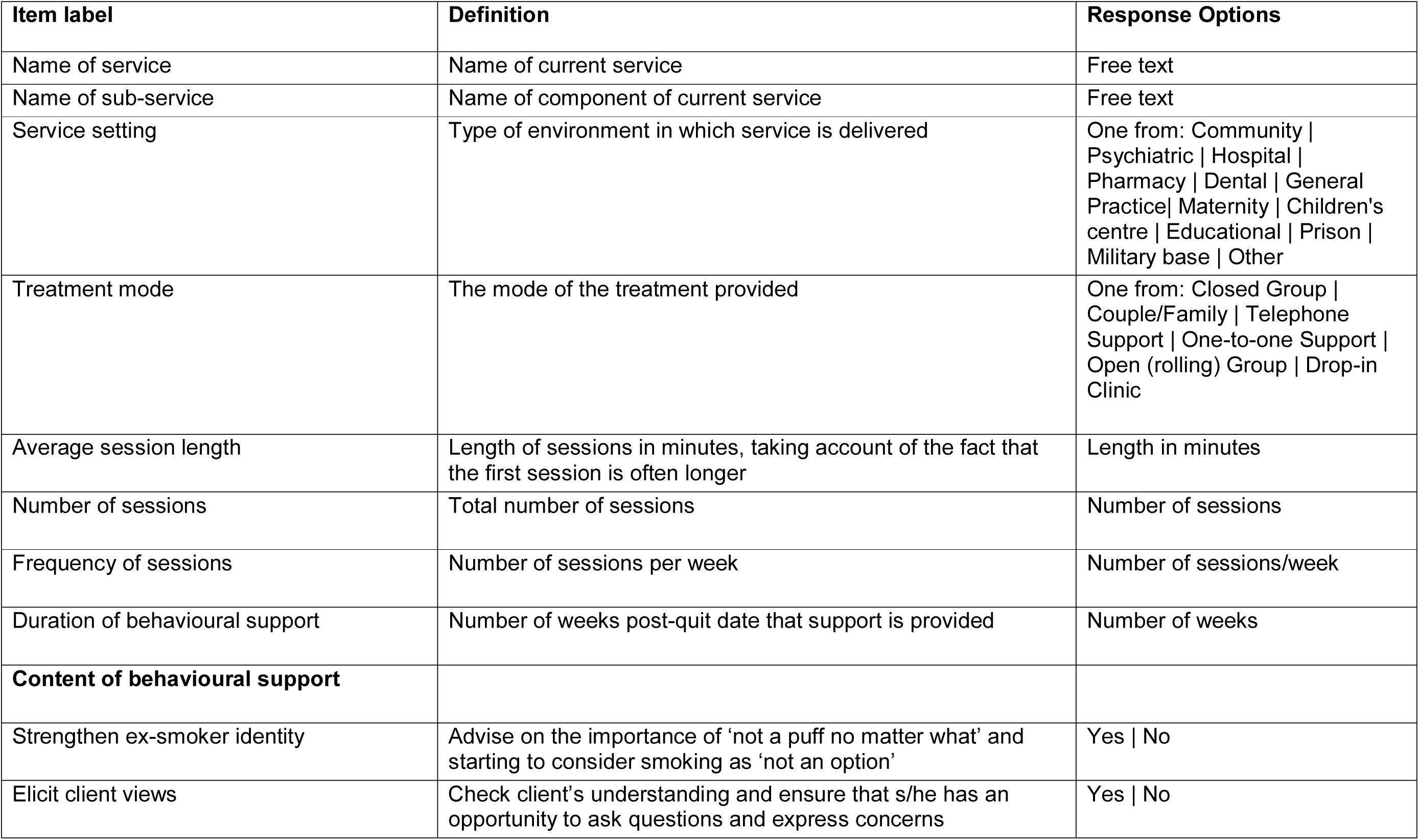
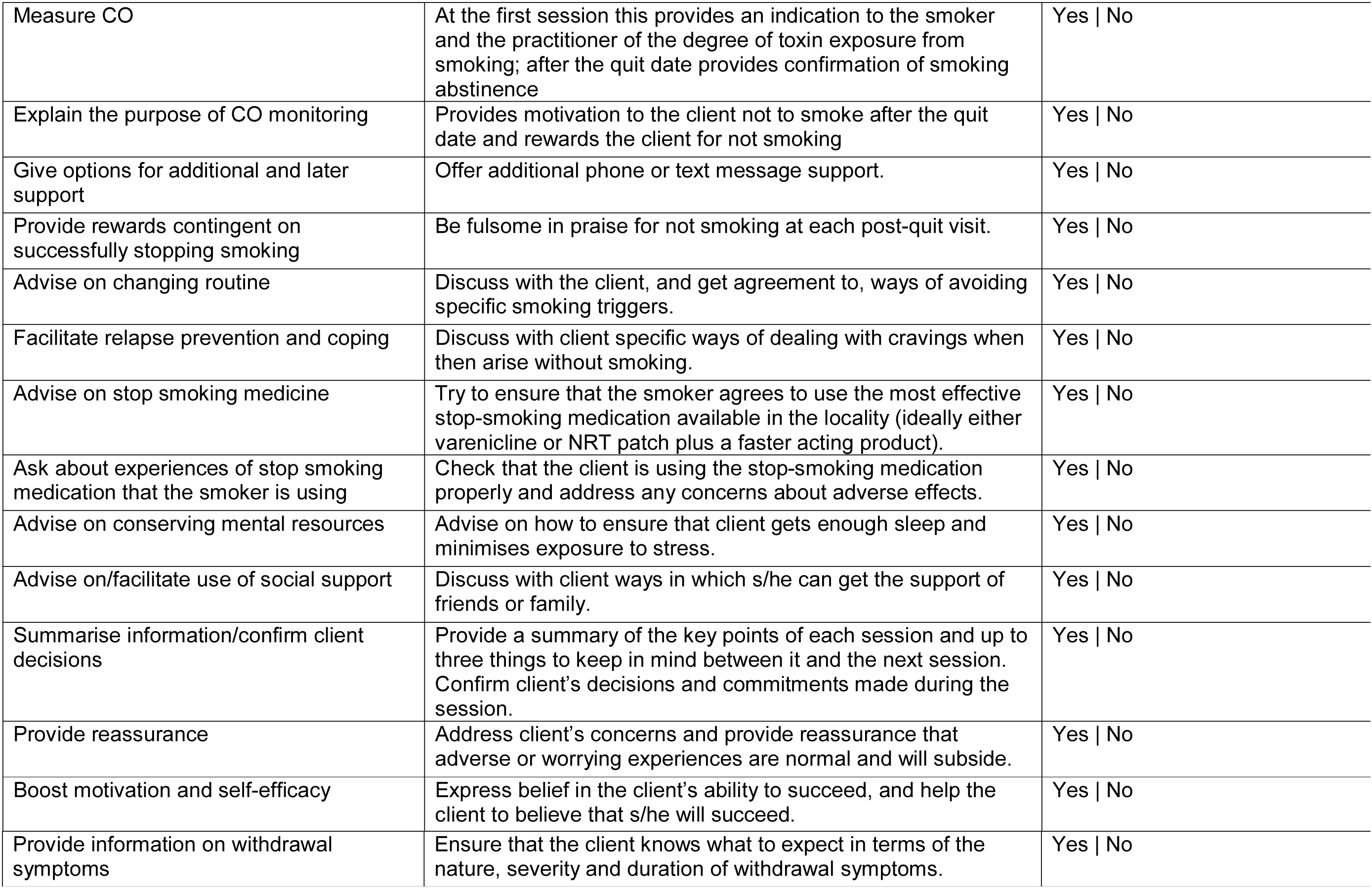
Treatment information (one record per type of treatment on offer)

| Item label | Definition | Response Options |
| --- | --- | --- |
| Name of service | Name of current service | Free text |
| Name of sub-service | Name of component of current service | Free text |
| Service setting | Type of environment in which service is delivered | One from: Community \| Psychiatric \| Hospital \| Pharmacy \| Dental \| General Practice \| Maternity \| Children's centre \| Educational \| Prison \| Military base \| Other |
| Treatment mode | The mode of the treatment provided | One from: Closed Group \| Couple/Family \| Telephone Support \| One-to-one Support \| Open (rolling) Group \| Drop-in Clinic |
| Average session length | Length of sessions in minutes, taking account of the fact that the first session is often longer | Length in minutes |
| Number of sessions | Total number of sessions | Number of sessions |
| Frequency of sessions | Number of sessions per week | Number of sessions/week |
| Duration of behavioural support | Number of weeks post-quit date that support is provided | Number of weeks |
| **Content of behavioural support** |  |  |
| Strengthen ex-smoker identity | Advise on the importance of 'not a puff no matter what' and starting to consider smoking as 'not an option' | Yes \| No |
| Elicit client views | Check client's understanding and ensure that s/he has an opportunity to ask questions and express concerns | Yes \| No |
| Measure CO | At the first session this provides an indication to the smoker and the practitioner of the degree of toxin exposure from smoking; after the quit date provides confirmation of smoking abstinence | Yes \| No |
| Explain the purpose of CO monitoring | Provides motivation to the client not to smoke after the quit date and rewards the client for not smoking | Yes \| No |
| Give options for additional and later support | Offer additional phone or text message support. | Yes \| No |
| Provide rewards contingent on successfully stopping smoking | Be fulsome in praise for not smoking at each post-quit visit. | Yes \| No |
| Advise on changing routine | Discuss with the client, and get agreement to, ways of avoiding specific smoking triggers. | Yes \| No |
| Facilitate relapse prevention and coping | Discuss with client specific ways of dealing with cravings when then arise without smoking. | Yes \| No |
| Advise on stop smoking medicine | Try to ensure that the smoker agrees to use the most effective stop-smoking medication available in the locality (ideally either varenicline or NRT patch plus a faster acting product). | Yes \| No |
| Ask about experiences of stop smoking medication that the smoker is using | Check that the client is using the stop-smoking medication properly and address any concerns about adverse effects. | Yes \| No |
| Advise on conserving mental resources | Advise on how to ensure that client gets enough sleep and minimises exposure to stress. | Yes \| No |
| Advise on/facilitate use of social support | Discuss with client ways in which s/he can get the support of friends or family. | Yes \| No |
| Summarise information/confirm client decisions | Provide a summary of the key points of each session and up to three things to keep in mind between it and the next session. Confirm client's decisions and commitments made during the session. | Yes \| No |
| Provide reassurance | Address client's concerns and provide reassurance that adverse or worrying experiences are normal and will subside. | Yes \| No |
| Boost motivation and self-efficacy | Express belief in the client's ability to succeed, and help the client to believe that s/he will succeed. | Yes \| No |
| Provide information on withdrawal symptoms | Ensure that the client knows what to expect in terms of the nature, severity and duration of withdrawal symptoms. | Yes \| No |

**Table 2 –.**
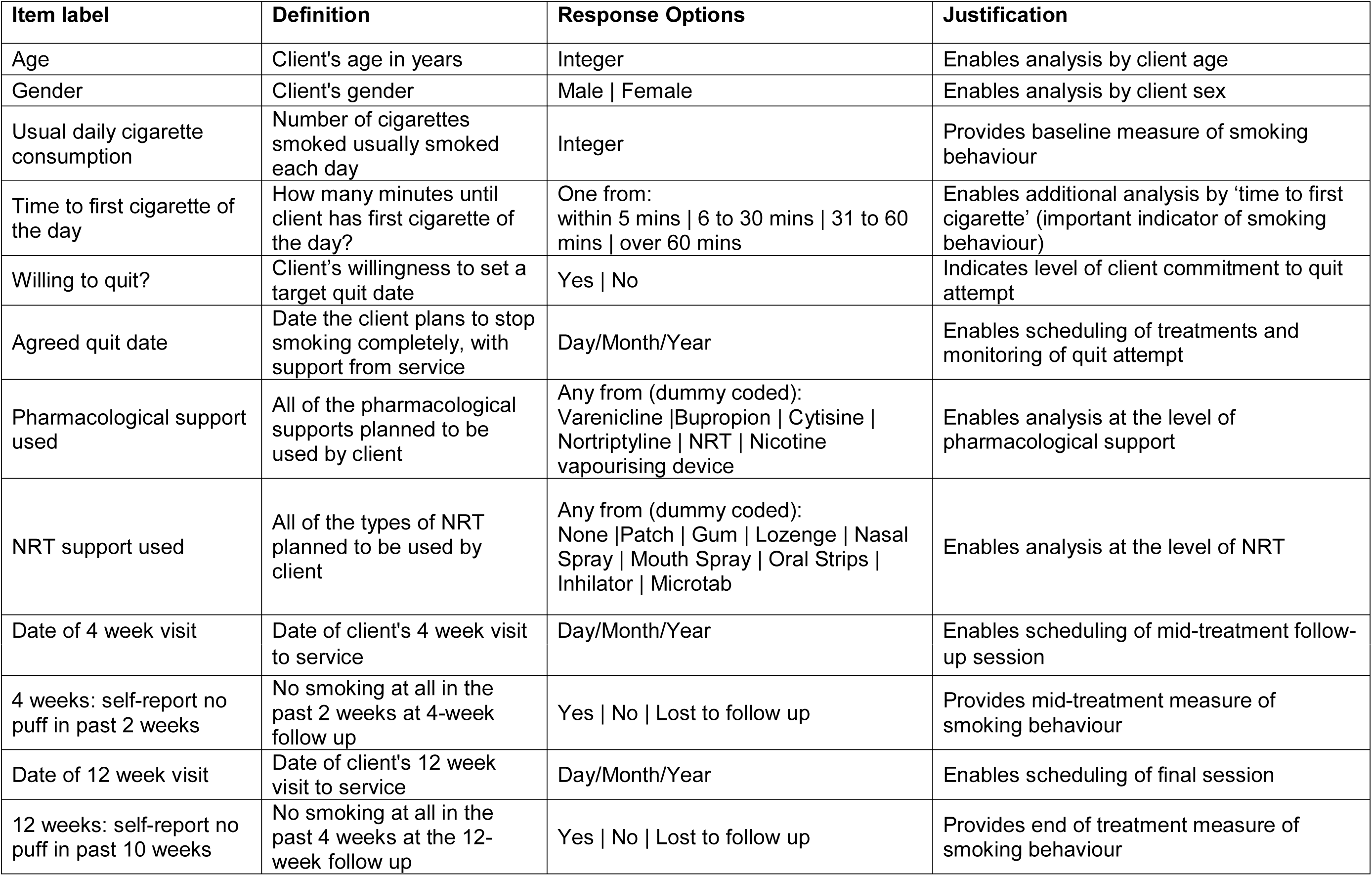
Core client data items

**Table 3 –.**
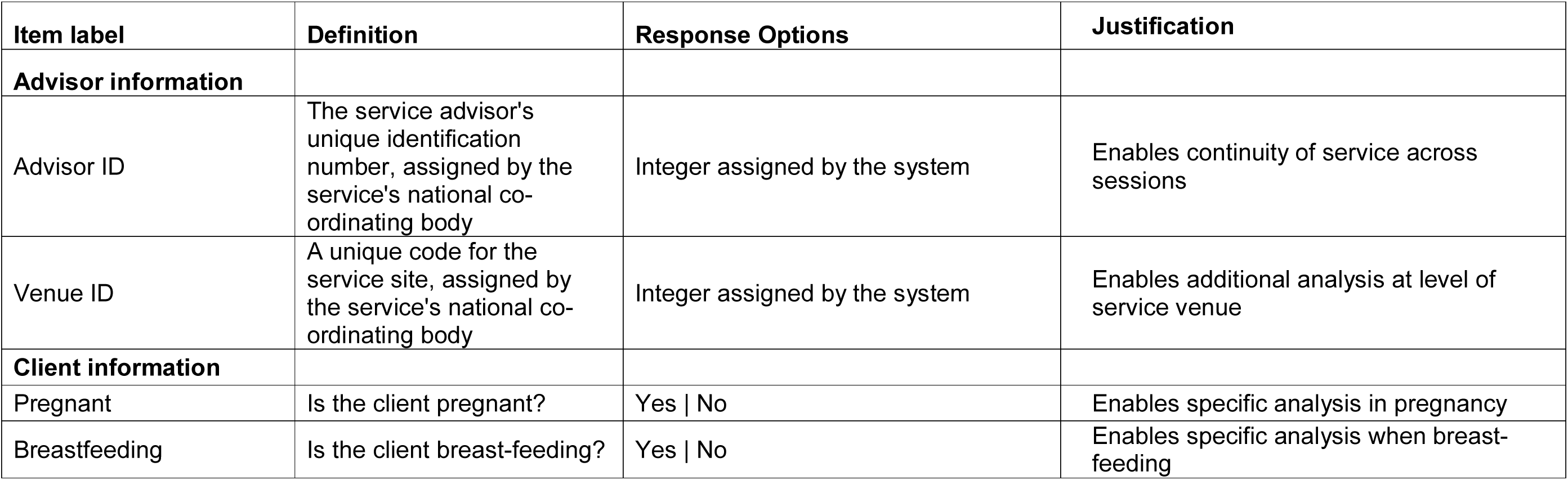
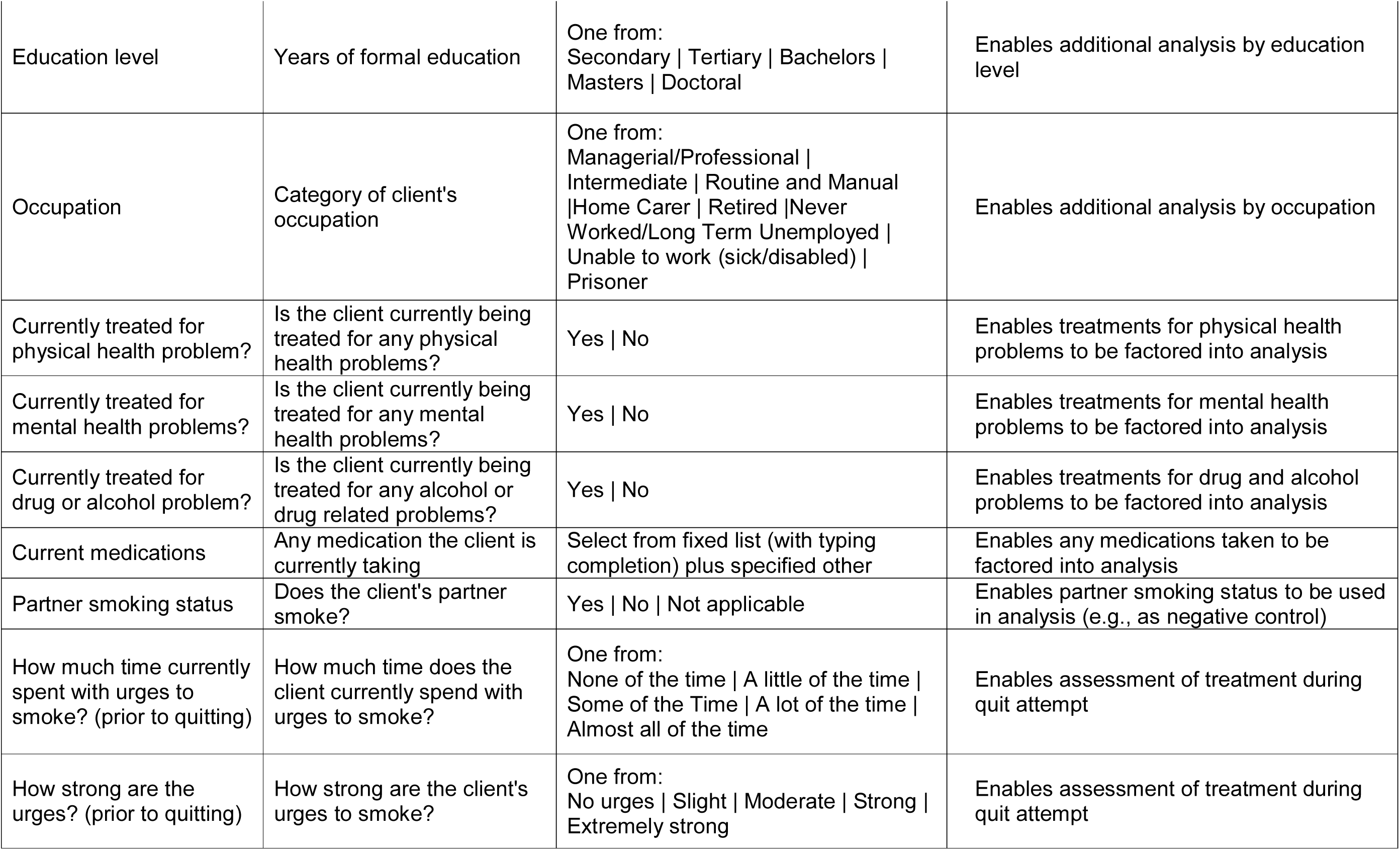
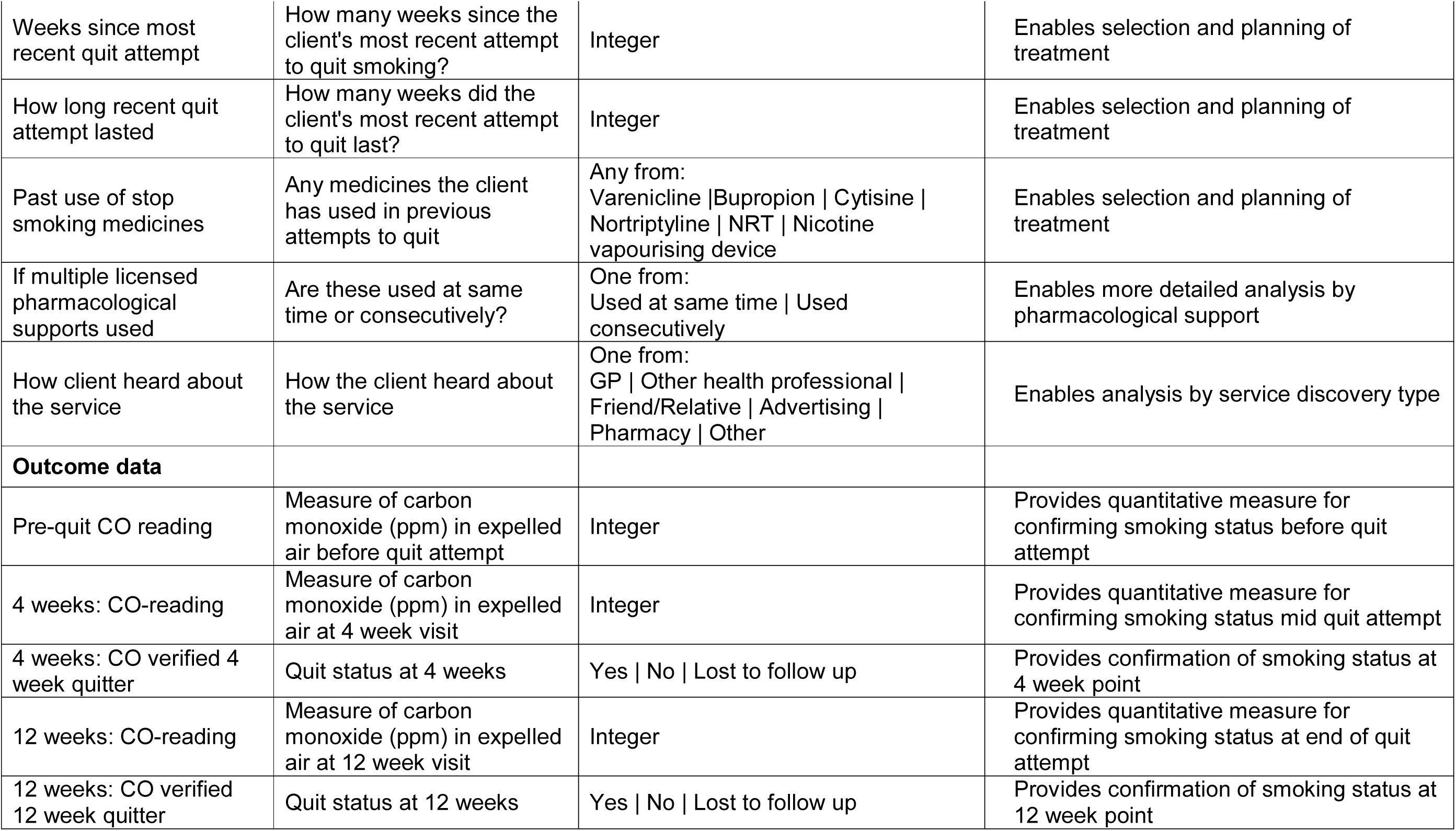
Extended client data items

| Item label | Definition | Response Options | Justification |
| --- | --- | --- | --- |
| **Advisor information** |  |  |  |
| Advisor ID | The service advisor's unique identification number, assigned by the service's national co-ordinating body | Integer assigned by the system | Enables continuity of service across sessions |
| Venue ID | A unique code for the service site, assigned by the service's national co-ordinating body | Integer assigned by the system | Enables additional analysis at level of service venue |
| **Client information** |  |  |  |
| Pregnant | Is the client pregnant? | Yes \| No | Enables specific analysis in pregnancy |
| Breastfeeding | Is the client breast-feeding? | Yes \| No | Enables specific analysis when breast-feeding |
| Education level | Years of formal education | One from:<br>Secondary \| Tertiary \| Bachelors \|<br>Masters \| Doctoral | Enables additional analysis by education level |
| Occupation | Category of client's occupation | One from:<br>Managerial/Professional \|<br>Intermediate \| Routine and Manual<br>\|Home Carer \| Retired \|Never<br>Worked/Long Term Unemployed \|<br>Unable to work (sick/disabled) \|<br>Prisoner | Enables additional analysis by occupation |
| Currently treated for physical health problem? | Is the client currently being treated for any physical health problems? | Yes \| No | Enables treatments for physical health problems to be factored into analysis |
| Currently treated for mental health problems? | Is the client currently being treated for any mental health problems? | Yes \| No | Enables treatments for mental health problems to be factored into analysis |
| Currently treated for drug or alcohol problem? | Is the client currently being treated for any alcohol or drug related problems? | Yes \| No | Enables treatments for drug and alcohol problems to be factored into analysis |
| Current medications | Any medication the client is currently taking | Select from fixed list (with typing completion) plus specified other | Enables any medications taken to be factored into analysis |
| Partner smoking status | Does the client's partner smoke? | Yes \| No \| Not applicable | Enables partner smoking status to be used in analysis (e.g., as negative control) |
| How much time currently spent with urges to smoke? (prior to quitting) | How much time does the client currently spend with urges to smoke? | One from:<br>None of the time \| A little of the time \|<br>Some of the Time \| A lot of the time \|<br>Almost all of the time | Enables assessment of treatment during quit attempt |
| How strong are the urges? (prior to quitting) | How strong are the client's urges to smoke? | One from:<br>No urges \| Slight \| Moderate \| Strong \|<br>Extremely strong | Enables assessment of treatment during quit attempt |
| Weeks since most recent quit attempt | How many weeks since the client's most recent attempt to quit smoking? | Integer | Enables selection and planning of treatment |
| How long recent quit attempt lasted | How many weeks did the client's most recent attempt to quit last? | Integer | Enables selection and planning of treatment |
| Past use of stop smoking medicines | Any medicines the client has used in previous attempts to quit | Any from:<br>Varenicline \| Bupropion \| Cytisine \| Nortriptyline \| NRT \| Nicotine vapourising device | Enables selection and planning of treatment |
| If multiple licensed pharmacological supports used | Are these used at same time or consecutively? | One from:<br>Used at same time \| Used consecutively | Enables more detailed analysis by pharmacological support |
| How client heard about the service | How the client heard about the service | One from:<br>GP \| Other health professional \| Friend/Relative \| Advertising \| Pharmacy \| Other | Enables analysis by service discovery type |
| **Outcome data** |  |  |  |
| Pre-quit CO reading | Measure of carbon monoxide (ppm) in expelled air before quit attempt | Integer | Provides quantitative measure for confirming smoking status before quit attempt |
| 4 weeks: CO-reading | Measure of carbon monoxide (ppm) in expelled air at 4 week visit | Integer | Provides quantitative measure for confirming smoking status mid quit attempt |
| 4 weeks: CO verified 4 week quitter | Quit status at 4 weeks | Yes \| No \| Lost to follow up | Provides confirmation of smoking status at 4 week point |
| 12 weeks: CO-reading | Measure of carbon monoxide (ppm) in expelled air at 12 week visit | Integer | Provides quantitative measure for confirming smoking status at end of quit attempt |
| 12 weeks: CO verified 12 week quitter | Quit status at 12 weeks | Yes \| No \| Lost to follow up | Provides confirmation of smoking status at 12 week point |

In addition to regional differences in resources available to capture data, there will also be regional variations in factors such as the way data items are labelled, the names of drugs, and the types of questions that are culturally acceptable to ask. To explore these issues, feedback on the initial proposals was sought from colleagues in the International Cooperation Centre for the Framework Convention on Tobacco Control in Uruguay, and the Centre for Tobacco Studies in Syria. In both cases the feedback was positive, with agreement that the choice of data items for all sets was appropriate. The main issue identified with the initial proposals was that they included CO measurements in the core client data set. The availability of CO measurement is limited in some regions, so CO readings and CO verification statuses were moved from the core to the extended client set. It was noted that ‘willingness to quit’ was not included, and this was added to the core client data set. Finally, it was noted that ‘gender’ would be more acceptable than ‘sex’, and that ‘age’ would be preferable to ‘date of birth’, and these changes were made.

## Results

The proposed set of treatment data items is shown in Table 1. The proposed set of core data items that should be captured in every clinical smoking cessation service setting is shown in Table 2. The proposed set of extended data items that should be captured where resources permit is shown in Table 3.

## Discussion

Whilst individual tobacco cessation clinics using locally-defined measures and methods may be effective to some degree, harmonising the data captured across clinics will make it possible to assess the effectiveness of different treatments in different regions, to identify optimal sets of approaches for supporting smoking cessation in a variety of different settings, and to commission more effective and cost-effective measures that can reach more smokers and increase quit rates. This approach has proved successful in the United Kingdom, through standardisation of outcome assessments, and use of common data structures in the systems supporting services. Our vision is to extend this approach internationally.

Here, we take the first steps by proposing sets of data items that should be captured by clinical smoking cessation services internationally. Our aim is that these proposals should be a starting point for a discussion that converges to identify optimal sets of data items. We invite comments and suggestions on any aspects of these, including; which items should be captured, whether they should be core or extended, and details about how they are labelled and response options. Once an agreed core set has been established it would be a simple matter to set up an online data service that can store the data and provide clinics, practitioners with on-demand access to performance information.

## Acknowledgements

The authors are members of the United Kingdom Centre for Tobacco and Alcohol Studies, a UKCRC Public Health Research: Centre of Excellence. Funding from British Heart Foundation, Cancer Research UK, Economic and Social Research Council, Medical Research Council (grants MC_UU_12013/6 and MC_UU_12013/7), and the National Institute for Health Research, under the auspices of the UK Clinical Research Collaboration, is gratefully acknowledged. The authors thank Prof Samer Rastam of the Syrian Center for Tobacco Studies and Amanda Sica of the Honorary Commission for the Fight Against Cancer, Uruguay for feedback on the proposed data sets.

